# SyntenyPair Explorer: an installation-free, browser-based tool for interactive pairwise genome synteny visualization

**DOI:** 10.64898/2026.07.23.740353

**Authors:** John G. Gibbons

## Abstract

Comparisons of genome structure between related organisms are central to understanding genome evolution, gene family dynamics, and the genomic basis of phenotypic variation. Synteny, the conserved co-localization of genes along chromosomes, is most readily interpreted visually, yet many existing synteny visualization tools require local software installation, command-line proficiency, and/or dedicated server infrastructure, and produce static images that cannot be explored interactively. Here, I present SyntenyPair Explorer, a lightweight, installation-free tool for interactive visualization of synteny between two genomes. The application runs entirely within a standard web browser as a single, self-contained HTML file with no external dependencies and no server-side component. SyntenyPair Explorer accepts standard file formats already produced by common comparative genomics workflows, including FASTA genome assemblies (used to compute optional assembly summary statistics), GFF3/GTF gene annotations, and either BLAST tabular output (outfmt 6) or MCScanX collinearity files. Syntenic relationships are resolved by gene-identifier matching between the relationship file and the gene annotations, so that each relationship corresponds to a discrete gene-to-gene link. Interactive features include continuous zoom and pan, gene search with automatic centering of the partner genome on the syntenic counterpart, synteny block coloring, extensively customizable gene highlights and annotation callouts, session saving and restoration, and publication-quality image export. I demonstrate the tool by visualizing structural differences at the alpha-amylase loci between two strains of the industrially important fungus *Aspergillus oryzae*. SyntenyPair Explorer lowers the technical barrier to interactive synteny visualization and is freely available under the MIT license at https://github.com/GibbonsLabGenomics/SyntenyPair-Explorer, with a live browser-based version at https://gibbonslabgenomics.github.io/SyntenyPair-Explorer/.

## INTRODUCTION

The comparison of genome organization across species and strains is a cornerstone of modern genomics. The relative order and orientation of genes along chromosomes, their synteny, carries important evolutionary information, reflecting the history of rearrangements, duplications, inversions, translocations, and losses that distinguish genomes [1]. Patterns of conserved synteny are routinely used to identify orthologous regions, trace the evolution of gene clusters, detect structural variation associated with adaptation or disease, transfer functional annotation between species, and reconstruct ancestral genome states [2–5]. As the number of sequenced and well-annotated genomes continues to grow rapidly across all domains of life, the ability to examine and communicate synteny relationships has become an increasingly routine practice for researchers who are not necessarily computational specialists.

Synteny is fundamentally a spatial relationship and is therefore effectively interpreted through visualization. A range of established tools support synteny analysis and display, each with particular strengths. MCScanX is widely used to detect collinear gene blocks and provides associated visualization utilities [6, 7]. The JCVI/MCscan Python toolkit [8] produces high-quality publication-ready synteny figures. Web-based platforms such as CoGe/SynMap [9, 10] offer powerful synteny analyses backed by server infrastructure. Circos produces visually striking circular representations of genomic relationships [11], and genome alignment browsers such as Mauve enable inspection of aligned regions [12]. For pairwise comparison of annotated regions specifically, tools such as Easyfig [13], genoPlotR [14], SynVisio [15], and the R packages gggenes/gggenomes [16] produce linear comparison figures of high quality.

Despite these numerous and powerful tools, a practical gap remains for researchers who want to interactively explore synteny between two genomes quickly and without technical overhead. Many existing tools require local installation and dependency management, familiarity with the command line or with the R or Python programming environments, or access to and upload of data to external servers. Others are oriented toward producing static, publication-ready figures rather than supporting the fluid, exploratory examination (e.g. zooming into a locus of interest, searching for a specific gene, following a syntenic block into its neighborhood in the other genome etc.) that often precedes figure generation and hypothesis formation.

Here, I present SyntenyPair Explorer, a tool designed to fill this gap by prioritizing accessibility, interactivity, and zero-friction deployment. SyntenyPair Explorer runs entirely in a web browser as a single self-contained HTML file. It requires no installation, no software dependencies, and no server: the executable html file can be opened directly from a local disk or accessed through a hosted link, and all computation occurs locally in the browser so that potentially sensitive or unpublished genomic data never leave the user’s machine. The tool accepts standard comparative genomics file formats (FASTA genome assemblies, GFF3 or GTF annotation files, and either BLAST tabular output or MCScanX collinearity files) and requires no reformatting or preprocessing. Syntenic relationships are resolved by matching gene identifiers between the relationship file and the gene annotations, so that each relationship corresponds to a discrete gene-to-gene link. For similarity searches, the aligned sequences are therefore supplied at the level of individual genes, coding sequences, or proteins rather than whole chromosomes. The interface emphasizes interactive exploration, with continuous zoom and pan, gene search that automatically locates and centers the syntenic counterpart in the partner genome, customizable synteny block coloring, flexible gene highlighting and annotation, session saving and restoration, and high-resolution image export for publication.

To illustrate the tool in a realistic research context, I use SyntenyPair Explorer to compare two strains of *Aspergillus oryzae*, a filamentous fungus of major importance to the food and biotechnology industries owing to its central role in the fermentation of sake, soy sauce, and miso, and its prolific secretion of hydrolytic enzymes [17]. The alpha-amylase loci of *A. oryzae*, which encode the secreted enzymes responsible for starch degradation during fermentation, are known to vary in copy number and genomic arrangement among strains [18–20], making them a biologically meaningful example of the structural variation that synteny visualization is well suited to reveal. This worked example demonstrates how SyntenyPair Explorer enables rapid, interactive inspection of locus-level structural differences between closely related genomes.

## MATERIALS AND METHODS

### Software development process

SyntenyPair Explorer was developed iteratively over approximately 200 interactive prompts across multiple extended sessions using Anthropic’s Claude (Sonnet 4.6), a large language model used as a code-generation tool under the direction and supervision of the author. The author conceived the tool, defined its requirements and target audience, and directed all design decisions throughout development. These included the choice of HTML5 Canvas rendering over SVG for scalability with large numbers of on-screen elements, the single-file zero-dependency architecture to eliminate installation barriers, and the gene-identifier-based link-resolution strategy with flexible identifier normalization. Over the course of development, more than 25 distinct features were implemented and iteratively refined. All functionality was tested and validated by the author on multiple datasets with different origins.

### Design goals and overall architecture

SyntenyPair Explorer was designed around three guiding principles: (1) minimal barrier to use, (2) fully interactive exploration, and (3) operation on the file formats commonly used in comparative and evolutionary genomics. These principles motivated the central architectural decision to implement the entire application as a single, self-contained HTML document that runs in any modern web browser. The file bundles its markup, styling, and program logic together and depends on no external libraries, frameworks, build steps, or network resources. Consequently, the tool can be used by opening a local copy of the file directly in a browser or by accessing a hosted copy through a URL (https://gibbonslabgenomics.github.io/SyntenyPair-Explorer/). Because all parsing, computation, and rendering occur locally within the browser, no genomic data are transmitted to any server.

The application is written in standard HTML, CSS, and vanilla JavaScript, and uses the HTML5 Canvas API for rendering. The Canvas API was chosen over scalable vector graphics (SVG) for the main visualization because the number of on-screen elements (i.e. gene symbols, synteny ribbons, axis ticks, and annotations etc.) can become large when whole contigs are displayed, and immediate-mode canvas drawing scales more gracefully than retaining thousands of individual document nodes. The interface is organized into a data-loading view and a visualization view, with the latter exposing interactive controls for navigation, styling, annotation, and export.

### Input formats and parsing

SyntenyPair Explorer requires two categories of input: gene annotations for each of the two genomes, and a file describing the syntenic or homologous relationships between them. Gene annotations are supplied as GFF3 or GTF files. The parser extracts gene-level features and accommodates the structural conventions of both formats, including the hierarchical relationship between gene, mRNA/transcript, and child features in GFF3. To avoid drawing the same locus multiple times when a gene is represented by several nested features, only root-level gene features are retained, identified by the absence of a parent attribute. Where gene-level features are not present, the parser falls back to transcript-level features and finally to positional deduplication. To accommodate the differing gene-identifier conventions used by common annotation sources, the parser matches identifiers flexibly, tolerating differences such as source-specific prefixes and version suffixes (e.g., matching gene-ID1G00001.1 to ID1G00001).

Syntenic relationships are supplied either as tabular sequence-similarity output in the standard “outfmt 6” format, as produced by BLAST or DIAMOND [21, 22], or as MCScanX collinearity files [6, 7] (the format is detected automatically from file content). This flexibility reflects the two most common ways in which pairwise relationships are generated in practice: (1) direct sequence-similarity search of one strain’s gene, coding-sequence, or protein set against another’s (e.g. BLAST or DIAMOND), and (2) dedicated collinearity analysis (e.g. MCScanX).

### Gene identifier–based link resolution

SyntenyPair Explorer resolves syntenic and homologous relationships by matching gene identifiers between the relationship file and the parsed gene annotations. Both supported relationship formats (MCScanX collinearity files and tabular BLAST or DIAMOND output in outfmt 6) reference the related sequences by gene identifier, so that each record corresponds to a single gene-to-gene relationship. For similarity searches, this means that the query and subject sequences should be individual genes, coding sequences, or proteins labeled by gene identifier (e.g., tabular output generated by searching one strain’s protein or coding-sequence set against another’s). For each record, the tool matches the query and subject identifiers against the annotations, applying the same identifier-normalization rules used during annotation parsing and testing both orientations of the pair. This gene-level matching resolves relationships precisely and avoids the ambiguity that can arise when alignments do not correspond to discrete genes. We note that raw whole-genome nucleotide alignments (e.g., genome vs. genome BLASTn), in which individual high-scoring segment pairs may span multiple genes, cover intergenic regions, or fragment a single gene across many alignment records, are not appropriate input for SyntenyPair Explorer, because such records cannot be unambiguously assigned to individual gene pairs. Comparison of whole assemblies should instead supply relationships at the level of genes, coding sequences, or proteins.

### Visualization and interaction

The two genomes are displayed as horizontal tracks, with the first genome above the second (**Figure 1A**). Genes are drawn as directional arrow glyphs whose orientation reflects the annotated strand and whose color encodes strand by default. Resolved syntenic relationships are drawn as shaded ribbons connecting the corresponding genes on the two tracks. Ribbons may be colored by collinear block to emphasize the modular structure of conserved synteny, or by percent identity where that information is available (i.e. from BLAST result input). Individual blocks can be toggled on and off to reduce visual complexity.

**Figure 1.**
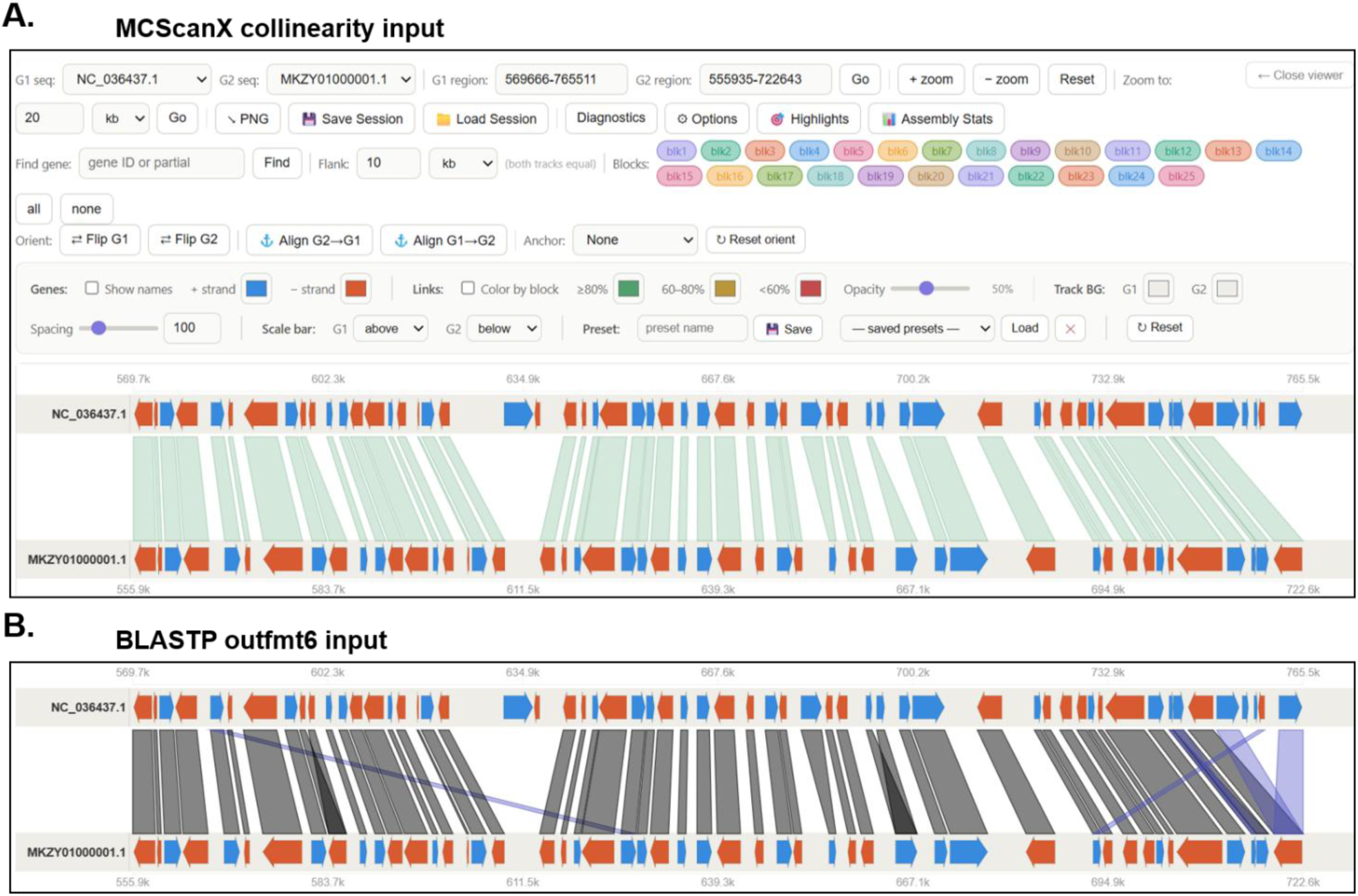
The SyntenyPair Explorer interface and the effect of relationship input on pairwise synteny visualization. A syntenic region between *Aspergillus oryzae* strain RIB40 (top track, chromosome NC_036437.1) and strain BCC7051 (bottom track, scaffold MKZY01000001.1) is shown. (A) Complete application view, comprising the data-loading and visualization controls (top) and the rendered comparison (bottom), using MCScanX collinearity output as the relationship input. Genes are drawn as strand-colored arrows (blue, plus strand; orange, minus strand) on each genome track, and shaded ribbons connect syntenic gene pairs. (B) The same genomic region rendered from tabular protein BLAST output (outfmt 6), generated by comparing the annotated protein sets of the two strains with BLAST+ v2.16.0 at an E-value cutoff of 1e-10. In both panels, relationships are resolved by matching gene identifiers between the relationship file and the gene annotations. In (B), ribbons are colored by percent amino-acid identity (grey, ≥80%; yellow, 60–80%; blue, <60%; no relationships in this region fall in the 60– 80% range), distinguishing the conserved syntenic backbone from more divergent relationships. Relative to the collinearity-based view in (A), the similarity-based view in (B) additionally recovers divergent and non-collinear relationships (blue ribbons), illustrating how the choice of relationship input can influence the interpretation of synteny.

Navigation is continuous and interactive. Users can zoom and pan each track with the mouse, specify exact coordinate ranges or a target window size, and search for a gene by identifier or annotation text. When a gene is located, the tool automatically positions the partner genome on the syntenic counterpart of the queried gene, optionally framing a defined number of flanking genes on either side, so that a locus and its conserved neighborhood can be examined in both genomes without manual coordinate lookup. Track orientation can be reversed independently for each genome to accommodate inversions and differing assembly conventions, and one track can be configured to follow the other during navigation.

To support figure preparation, SyntenyPair Explorer provides an adaptable gene-highlighting system. Sets of genes can be highlighted by identifier and styled with custom arrow, body, pointer, and label colors. Highlighted genes may be annotated with callout labels drawn from the gene’s functional annotation, its identifier, or user-supplied text. Additionally, individual labels can also be edited directly on the visualization. Callout labels are placed using a rotation-aware layout algorithm that computes the horizontal footprint of each label at its chosen angle and assigns labels to stacked rows so that they do not overlap, with the drawing area expanding automatically to accommodate labels of arbitrary length and orientation. The complete state of a session, including loaded data, view positions, styling, and annotations, can be saved to a file and later restored, and visual styling presets can be stored in the browser for reuse. Additionally, session settings presets (e.g. color coding, spacing, etc.) can be saved and loaded in subsequent analysis to aid in consistency during figure generation. Finished visualizations can be exported as high-resolution PNG images suitable for publication.

### Synteny analysis of *Aspergillus oryzae* RIB40 and BCC7051

To demonstrate the tool, genome assemblies for *A. oryzae* strains RIB40 and BCC7051 were obtained from the NCBI genomes database (RefSeq accession GCF_000184455.2 and GenBank accession GCA_002007945.1 for RIB40 and BCC7051, respectively) [23, 24]. To limit bias caused by predicted genes from different methods, I used AUGUSTUS v3.5.0 for gene prediction in both genomes using the following command line arguments: species=aspergillus_oryzae, --strand=both --genemodel=complete, --gff3=on, --codingseq=on, and --protein=on [25]. Strain names were appended to the gene identifiers to ensure unique gene IDs in the coding sequence and protein files, and the gff3 file.

For the collinearity-based analysis, MCScanX was used to identify collinear gene blocks between the two strains. MCScanX requires two inputs: a tabular all-versus-all protein similarity file and a combined gene-position file listing the chromosomal location of every gene in both genomes. The protein sequences of both strains were concatenated into a single FASTA file, and an all-versus-all protein comparison was performed with DIAMOND v2.1.10 [22] in “very-sensitive” mode (diamond blastp), retaining tabular output (outfmt 6) with an E-value cutoff of 1e-10, and up to five target sequences per query. The combined gene-position file listed, for each gene, includes its chromosomal/scaffold sequence identifier, a strain-prefixed gene identifier (e.g., AO_RIB40|g1, AO_BCC7051|g1), and its start and end coordinates, with genes from both strains merged into a single file. MCScanX was then run on these inputs with default parameters to produce the collinearity file used as input to SyntenyPair Explorer. For the similarity-based analysis, the annotated protein sets of the two strains were compared by all-versus-all protein BLAST (BLAST+ v2.16.0 [26]) with an E-value cutoff of 1e-10, retaining tabular output (outfmt 6).

The resulting annotation and relationship files (MCScanX collinearity and BLASTP outfmt6) were loaded directly into SyntenyPair Explorer, and the alpha-amylase loci were located using the built-in gene search (RIB40 = AO_RIB40|g3609, AO_RIB40|g4692 and AO_RIB40|g7934, and BCC7051 = AO_BCC7051|g4743 and AO_BCC7051|g5882). All figures in the worked example were exported directly from the tool. The example datasets are provided in the SyntenyPair Explorer GitHub software repository (https://github.com/GibbonsLabGenomics/SyntenyPair-Explorer/tree/main/example_data).

### Assembly statistics

When genome assembly files in FASTA format are provided as optional input alongside the required annotation files, SyntenyPair Explorer parses each assembly and computes a set of standard summary statistics. For each sequence (chromosome, scaffold, or contig), the tool reports its length, GC content, and the proportion of ambiguous (N) bases. At the assembly level, the tool reports the total number of sequences, total assembly length, mean and median sequence length, the longest and shortest sequences, and the N50, L50, N90, and L90 genome assembly contiguity metrics. These statistics are displayed in a toggle panel within the viewer interface and can be exported as a tab-separated file for use in downstream analyses or supplementary materials. Because the FASTA input is optional, the core synteny visualization functions identically without it. The assembly statistics panel simply provides additional context about the two genomes being compared.

## RESULTS

### Worked Example

To demonstrate the tool, genome assemblies for *Aspergillus oryzae* strains RIB40 and BCC7051 were compared [23, 24]. Pairwise relationships between the two strains were generated in two ways. First, collinear gene blocks were identified with MCScanX. Second, the annotated protein sets of the two strains were compared by all-versus-all protein BLAST [26]. The resulting annotation and relationship files were loaded directly into SyntenyPair Explorer, and the alpha-amylase loci were located using the built-in gene search.

### Overview and interface

SyntenyPair Explorer presents a single-page interface in which the two genomes are displayed as horizontal tracks and their syntenic relationships as connecting ribbons. All navigation, styling, annotation, and export controls are exposed directly in the browser, and the visualization updates interactively as the user zooms, pans, searches for genes, or adjusts display options. To illustrate both the interface and a key practical consideration in synteny visualization, **Figure 1** shows a representative syntenic region between *A. oryzae* strains RIB40 and BCC7051 rendered from two different relationship inputs.

Because SyntenyPair Explorer resolves relationships by gene-identifier matching, it can render synteny from either dedicated collinearity analysis or direct sequence-similarity search, and the choice between these inputs can affect the resulting visualization. Using MCScanX collinearity output (**Figure 1A**), the region is spanned by a set of collinear ribbons reflecting the conserved gene order between the two strains. Rendering the same region from protein BLAST output (**Figure 1B**) reproduces this conserved backbone but additionally recovers relationships between more divergent or non-collinear genes, shown here colored by amino-acid identity. This difference reflects the underlying methods: MCScanX enforces collinearity constraints when defining blocks, whereas direct similarity search reports all significant hits, including those between homologous or rearranged genes.

### Assembly summary

Loading the optional FASTA genome assemblies alongside the annotation files enabled SyntenyPair Explorer to compute and display assembly-level statistics for both strains. The RIB40 reference assembly comprised 12 sequences totaling 37,912,014 bp with a GC content of 47.2% and 2.10% ambiguous bases, an N50 of 4,887,096 bp, and scaffold lengths ranging from 6,938 to 6,520,266 bp. The BCC7051 assembly comprised 25 sequences totaling 38,507,590 bp with a GC content of 47.2% and no ambiguous bases, an N50 of 4,278,514 bp, and scaffold lengths ranging from 1,886 to 5,043,372 bp. These statistics are accessible through the Assembly Stats panel in the viewer and provide immediate context for interpreting the synteny visualization.

### Worked example: alpha-amylase copy-number variation between *A. oryzae* strains

To illustrate how SyntenyPair Explorer supports biological interpretation, I used it to examine the copy number variable alpha-amylase loci of *A. oryzae*, which encode the secreted enzyme central to starch degradation during traditional and industrial fermentation [18–20]. *A. oryzae* RIB40 carries three alpha-amylase gene copies, whereas strain BCC7051 carries two (**Figure 2A**). Using the built-in gene search to locate each copy and its syntenic context, SyntenyPair Explorer resolves this copy-number difference into its individual components and shows, for each copy, whether it is conserved or strain-specific (**Figure 2**).

**Figure 2.**
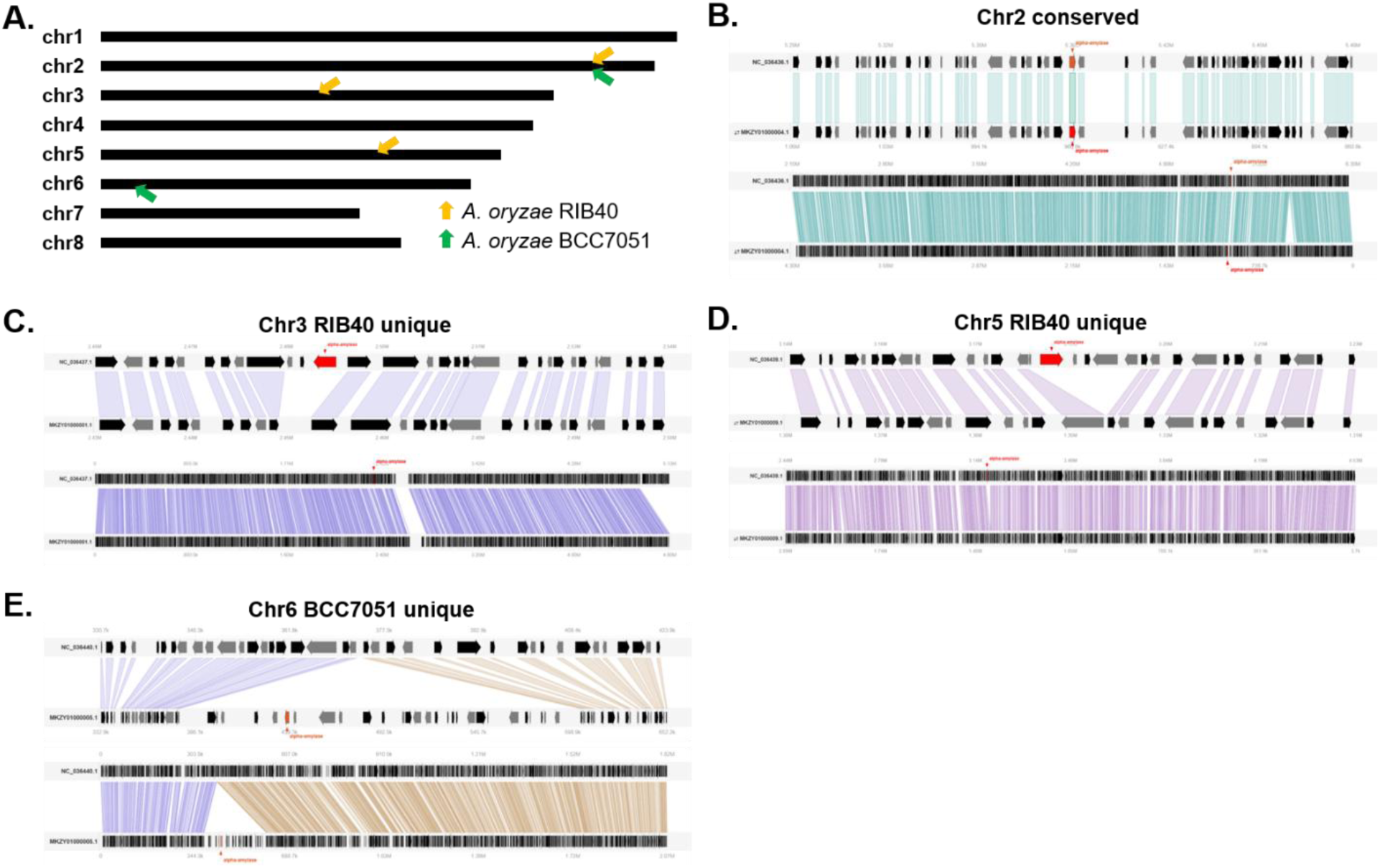
SyntenyPair Explorer resolves alpha-amylase copy-number variation between *Aspergillus oryzae* strains RIB40 and BCC7051 into conserved and strain-specific loci. All comparisons show strain RIB40 against strain BCC7051, visualized directly in the browser using MCScanX collinearity input. **(A)** Genomic positions of alpha-amylase gene copies across the eight RIB40 chromosomes. Arrows mark copies present in RIB40 (yellow) and BCC7051 (green). RIB40 carries three alpha-amylase copies and BCC7051 carries two. In panels B–E, each locus is shown at a gene-level zoom (top) and at a whole-chromosome or whole-contig zoom (bottom), and the alpha-amylase gene is highlighted in red. **(B)** The alpha-amylase copy on chromosome 2 is conserved between the strains and within a syntenic block. **(C, D)** Two alpha-amylase copies present in RIB40 lack a syntenic counterpart in BCC7051 (RIB40 unique). **(E)** An alpha-amylase copy present in BCC7051 that is absent from the syntenic position in RIB40 (BCC7051 unique). Genes are drawn as strand-colored arrows and shaded ribbons connect syntenic gene pairs. Track labels indicate the RIB40 chromosome and BCC7051 scaffold.

An overview of the genomic positions of all alpha-amylase copies across the eight RIB40 chromosomes (**Figure 2A**) shows that the copies are distributed across multiple chromosomes and establishes which copies are shared between the two strains and which are unique to one. Examining each locus individually clarifies the underlying pattern. One copy, located on chromosome 2, lies within a syntenic block conserved between the two strains, with clear one-to-one synteny of the alpha-amylase gene and its flanking neighborhood (**Figure 2B**). Two further copies are present in RIB40 but lack a syntenic counterpart in BCC7051 (**Figure 2C, D**). At these loci, the alpha-amylase gene and its immediate neighborhood in RIB40 have no corresponding syntenic region in BCC7051. Conversely, one copy is present in BCC7051 but absent from the syntenic position in RIB40 (**Figure 2E**). Displaying each locus at both a gene-level zoom and a whole-chromosome or whole-contig zoom (**Figure 2B–E**) allows the local structure of each alpha-amylase copy and its position within the broader conserved synteny to be examined together. These views resolve the copy-number difference into a locus-by-locus account of which copies are conserved and which are strain-specific, turning a summary count comparison into a deeper genome evolution insight.

## DISCUSSION

The worked example illustrates how interactive synteny visualization can move beyond summary statistics to provide a mechanistic, locus-level view of genome structural variation. Alpha-amylase copy-number variation among *A. oryzae* strains has been documented previously [18–20] and is thought to reflect selection during domestication for enhanced starch-degrading capacity [17, 20]. However, a simple count of gene copies does not distinguish between copies that are positionally conserved and those that are strain-specific, nor does it reveal whether strain-specific copies occupy syntenic positions that are otherwise conserved or represent genuinely novel insertions. By visualizing each locus in its syntenic context, SyntenyPair Explorer makes these distinctions immediately apparent and enables researchers to formulate specific hypotheses about the mechanisms underlying copy-number change, such as tandem duplication, segmental duplication, or non-homologous insertion [27]. This type of locus-level interrogation is applicable to any gene family or structural variant of interest across any pair of annotated genomes, and is not limited to fungal systems.

SyntenyPair Explorer is subject to several limitations that define its intended scope. The tool visualizes exactly two genomes and does not support multi-genome comparisons, whole-genome dot plots, or circular layouts, which are better served by existing tools such as Circos [11], SynVisio [15], or the JCVI toolkit [8]. Because syntenic relationships are resolved exclusively by gene-identifier matching, the tool requires that input relationship files reference individual genes, coding sequences, or proteins by identifier. Raw whole-genome nucleotide alignments, in which alignment coordinates do not correspond to discrete genes, are not supported. The quality of the visualization is therefore dependent on the quality of the underlying gene annotation and the relationship-inference method used [28]. Additionally, because all data are parsed and rendered in the browser, performance may degrade for very large genomes with tens of thousands of genes per chromosome, although the tool performs well for fungal-scale genomes (∼30–40 Mb) as demonstrated here. Finally, the current implementation does not compute syntenic relationships itself. It consumes the output of dedicated tools such as MCScanX or BLAST/DIAMOND, which must be run separately.

With that being said, SyntenyPair Explorer provides an interactive, installation-free environment for visualizing synteny between two genomes. By running entirely in the browser from a single self-contained file, operating on file formats that comparative genomics workflows already produce, and processing all data locally, it lowers the technical barrier to interactive synteny visualization for researchers who are not computational specialists while keeping potentially sensitive data on the user’s own machine. These properties distinguish SyntenyPair Explorer from the existing collection of synteny and genome-comparison tools, each of which excels in a particular niche. Command-line packages such as MCScanX [6] and the JCVI/MCscan toolkit [8] provide powerful collinearity detection and high-quality figure generation, but require local installation, dependency management, and familiarity with the command line, and their outputs are primarily static. Circos [11] produces striking circular genome-relationship figures, and Mauve [12] enables inspection of whole-genome alignments, but both are oriented toward configured or precomputed views rather than lightweight, on-the-fly exploration of a locus of interest. Web-based platforms such as CoGe/SynMap [9] offer sophisticated synteny analyses, but depend on server infrastructure and require uploading data to a remote service, which is not always desirable for unpublished genomes. Dedicated linear pairwise-comparison tools, including Easyfig [13], genoPlotR [14], SynVisio [15], and the R packages gggenes and gggenomes [16], produce excellent publication figures, but variously require local installation, programming in R, or a prior analysis pipeline, and are generally designed to render a defined figure rather than to support fluid, interactive navigation. SyntenyPair Explorer is not intended to replace these tools, several of which perform analyses (such as collinear block detection) that it consumes as input. Rather, SyntenyPair Explorer complements them by providing an accessible, interactive front end for examining pairwise synteny with no installation, no programming, and no server. Its most direct contribution is to make the exploratory phase of synteny analysis (e.g. searching for a gene, examining its conserved neighborhood, comparing the effect of different relationship inputs, and preparing an annotated figure) fast and approachable for a broad range of users.

Future planned development includes support for visualization of more than two genomes, additional relationship-file formats, GC-content visualization from FASTA input, SVG export for vector-editable figures etc. Community contributions are welcomed through the project’s public repository. By prioritizing accessibility and interactivity, SyntenyPair Explorer aims to make routine synteny visualization a low-friction part of routine comparative genomics.

## AVAILABILITY

SyntenyPair Explorer is freely available under the MIT License. Source code and documentation are hosted at https://github.com/GibbonsLabGenomics/SyntenyPair-Explorer. A live, ready-to-use version runs in the browser at https://gibbonslabgenomics.github.io/SyntenyPair-Explorer/.

A permanently archived version of the release described here is deposited at Zenodo (DOI: https://doi.org/10.5281/zenodo.20646099). The *Aspergillus oryzae* example dataset used in the worked example is provided in the repository.

## ACKNOWLEDGEMENTS

This work was supported by the National Science Foundation grant 1942681 to J.G.G.

## AI USAGE DISCLOSURE

Anthropic’s Claude (Sonnet 4.6) was used as an interactive code-generation and drafting tool during the development of SyntenyPair Explorer and in the preparation of this manuscript. All scientific decisions, software design, testing, validation, and final editorial judgment were performed by the author.

